# Slow Wave Entrainment Using a Smartwatch: A Randomized Crossover Study

**DOI:** 10.1101/2025.07.28.666762

**Authors:** Nathan W Whitmore, Samantha W.T. Chan, Abigail Dulski, Anita Podrug, Nelson Hidalgo, Ella F. Tubbs, Nnamdi Obi, Varun K. Viswanath, Michael S. Freedman, Viswam Nathan, Pattie Maes

**Author notes:** Equal contributions (Samantha W.T. Chan, Abigail Dulski). **Corresponding author:** Nathan W. Whitmore, 75 Amherst St, Cambridge, MA, 02139.

## Abstract

**Background:** Slow-wave sleep is critical for sleep quality, cognitive function, and mood. Slow-wave entrainment (SWE) via rhythmic sensory stimulation can enhance slow-wave activity. However, existing implementations rely on EEG systems, thereby limiting accessibility and scalability. Consumer smartwatches offer an opportunity to deliver SWE in home settings without EEG hardware.

**Objective:** This study evaluated whether smartwatch-delivered sensory stimulation applied during smartwatch-estimated deep sleep elicits acute changes in frontal slow-wave EEG activity during home sleep, and whether individual differences in neural responsiveness to stimulation are associated with next-day behavioral and sleep measures.

**Methods:** In a randomized crossover design, participants recruited offline from the Boston area slept at home for two nights while wearing a consumer smartwatch for stimulation delivery and a portable EEG headband for neural recording. On a single night, participants received block-wise auditory, vibrotactile, or combined stimulation, guided by an automated on-watch sleep-staging model based on heart rate and motion. On the other night, no stimulation was delivered. Event-related changes in frontal delta (1–4 Hz) power were quantified relative to pre-stimulation baselines. Sleep disruption, subjective sleep quality, mood, and cognitive performance were assessed using questionnaires and a computerized Trail Making Test emailed to participants and completed online.

**Results:** Initiation of sensory stimulation was associated with significant increases in frontal delta power relative to pre-stimulation baseline and matched non-stimulation blocks.Stimulation blocks exhibited lower disruption rates than non-stimulation blocks, suggesting improved sleep stability during stimulation periods. No significant group-level differences were observed between stimulation and non-stimulation nights on measures of sleep quality, mood, or cognition. However, across participants, larger stimulation-evoked increases in delta power were associated with more favorable next-day subjective sleep and mood ratings and fewer clicks to complete the Trail Making Test. 68/93 participants were stimulated overnight.

**Conclusions:** Smartwatch-based slow-wave entrainment delivered during home sleep can elicit reproducible delta EEG responses without sleep disruption. Individual differences in neural responsiveness to stimulation were associated with next-day behavioral measures, suggesting that wearable-based SWE may represent a scalable and accessible approach for improving sleep health.

**Trial Registration:** The experiment was retrospectively registered at ISRCTN (registration number pending)

## Introduction

The Centers for Disease Control and Prevention (CDC) reports that sleep-related problems affect 50-70 million people in the United States, with a large proportion of adults reporting insufficient or poor-quality sleep[1]. Poor sleep quality is associated with impairments in mood, attention, and memory, and, over long periods, increased risk of adverse cardiovascular, cardiometabolic, and mental health outcomes[1–3]. Poor sleep quality can be secondary to other conditions, such as sleep apnea[4], lifestyle and environmental factors[5,6], or idiopathic causes. Sleep quality also shows a bidirectional relationship with mental health[7], aging, and neurodegenerative disease[8], whereby sleep disruption both accompanies and exacerbates disease progression.

Typical approaches to address poor sleep include treating contributing conditions, behavioral and lifestyle interventions, and pharmacological treatments when insomnia symptoms are present. While behavioral interventions can be effective, they are often difficult for patients to implement consistently due to lifestyle constraints[9]. Pharmacologic treatments are not recommended as first-line interventions for chronic insomnia because potential adverse effects may outweigh benefits for long-term sleep quality[10]. These limitations motivate continued exploration of alternative approaches to modulating sleep.

One promising approach to improving sleep quality is to modulate slow-wave activity during sleep. Slow-wave sleep (N3) is a deep phase of non-rapid eye movement (NREM) sleep characterized by prominent low-frequency oscillations, typically in the range of 0.5-1.5 Hz. The intensity of slow-wave activity during NREM sleep reflects homeostatic sleep need[11], and slow-wave neural activity has been associated with many restorative functions of sleep, such as memory, cognitive function, and mental health[3,12–15].

Evidence for the functional relevance of slow-wave activity has motivated the development of slow-wave entrainment (SWE) approaches, where external sensory stimulation is used to evoke or amplify slow-wave EEG responses during sleep. Prior studies have shown that SWE can increase the restorative benefits of sleep in multiple domains, including cognitive, immune, and cardiovascular function[16]. The most commonly studied form of SWE is closed-loop auditory stimulation (CLAS), which delivers brief auditory pulses phase-locked to naturally occurring slow waves detected using EEG recordings. CLAS is thought to increase the amplitude of slow waves through resonance-like interactions between sensory-evoked potentials, K complexes, and ongoing slow oscillations[17,18]. Although auditory stimulation itself is noninvasive and well tolerated, CLAS requires continuous EEG acquisition and real-time signal processing to detect sleep stage and slow-wave phase.

The requirement for EEG monitoring represents a practical limitation for widespread adoption of CLAS outside laboratory environments. While home EEG devices have been developed, their use remains constrained by factors such as user comfort, signal quality, and cost, limiting their accessibility for large-scale or long-term use[19–21].

Wrist-worn smartwatches that estimate sleep state and deliver sensory stimulation present a potentially more accessible and comfortable platform for sleep-related interventions. Devices such as the Samsung Galaxy Watch, Fitbit, and the Apple Watch have been adopted by nearly 44% of Americans, and algorithms using motion and heart rate signals can estimate sleep stages, albeit with lower accuracy than EEG-based methods[22]. Although such systems cannot phase-lock stimulation to individual slow waves, prior work suggests that timing-guided, non–phase-locked stimulation can elicit slow-wave EEG responses when delivered with appropriate intensity and timing to avoid disrupting sleep[23].

In this study, we tested whether SWE delivered via a Samsung Galaxy Watch 6 could elicit acute changes in slow-frequency EEG activity during home sleep. In particular we sought to address four questions: (1) Does smartwatch-SWE increase delta activity? (2) Does smartwatch SWE also produce improvements in subjective sleep, mood, and cognition the next day? (3) Does slow wave entrainment cause sleep disruption, and (4) What modalities and patterns are least disruptive and most effective? In the study, participants received one night with SWE and one night with no stimulation, with the SWE night additionally divided into blocks of different stimulation types which allowed us to compare stimulation types within-night.

Participants received rhythmic auditory and/or vibrotactile stimulation during smartwatch-estimated deep sleep while wearing a portable EEG headband for neural recording. Using a randomized crossover design, we assessed event-related changes in delta and slow oscillation power following stimulation onset and explored whether individual differences in stimulation-evoked neural responses were associated with next-day measures of subjective sleep quality, mood, and cognitive performance.

## Methods

### Participants

The study was conducted between April 8 and October 25 2024, with a total of 93 participants (27 male, 66 female) ranging in age from 18-92 years old (mean=43, median=38, SEM=2). Participants were recruited from the Boston/Cambridge community and also from the Laselle Village retirement community. As the study involved within subject/within night comparisons we did not systematically inquire about concomitant sleep interventions. Eligibility criteria were age between 18-120. The study was approved by the MIT IRB (COUHES protocol number 2309001115) and all participants gave informed consent at their first lab visit at the beginning of the study. Clinical trial number: pending.. Participants were instructed to use the watch and EEG device while sleeping on both nights. Because participants recorded data in their own homes, and were not always successful at doing so (Figure 7 shows a participant flow), not all participants had all types of data available. However we found that the subgroups of participants with specific data available had similar demographics to the overall sample (Table S1). We recruited participants until we reached 35 with good delta EEG data, a threshold selected based on practical considerations as well as general consistency with prior slow wave entrainment studies.

### Experimental Design

The study was structured as an exploratory crossover trial with the overall experimental design shown in Figure 1. Participants visited the laboratory on day 1 when they collected the study equipment, completed consent forms, and were oriented to the devices used. On day 1, participants also completed their initial surveys, consisting of the Pittsburgh Sleep Quality Index (PSQI)[24] and the General Health Questionnaire (GHQ-12)[25], as well as questions about their age and gender. Night 1 and night 2 were experimental nights; participants were randomly assigned to receive slow wave entrainment on either night 1 or night 2. On each night, participants donned the Muse EEG device and the Galaxy watch and began recording EEG data before going to sleep. On either night 1 or night 2 (randomized), stimulation was delivered in blocks of specific stimulus types as shown in Figure 1B. Following both night 1 and night 2, participants were emailed the post-sleep questionnaire. On this questionnaire, participants rated their sleep quality the previous night (Leeds Sleep Evaluation Questionnaire (LSEQ)[26]), current mood (Brunel Mood Scale (BRUMS)[27]), and performed a modified computerized version of the Trail Making Task B[28] (Figure 1C) to measure overall cognitive function. After completing night 2, participants returned the equipment to the lab. While participants slept, our on-watch sleep staging model performed sleep staging once per second for the duration of the whole night. Slow wave stimulation was delivered on stimulation nights when our custom on-watch sleep staging model determined the participant was likely in stage N3 sleep. The watch then began the stimulation protocol. We tested three distinct types of stimulation: stimulation with 50ms pink noise pulses, stimulation with 50ms vibration pulses, and stimulation with combined sound and vibration. Additionally, we tested two stimulation duty cycles: A “continuous” mode where stimuli were presented every 1.2 seconds, and a “short-train” mode where two stimuli were presented 1.2 seconds apart every 10 seconds.

**Figure 1:**
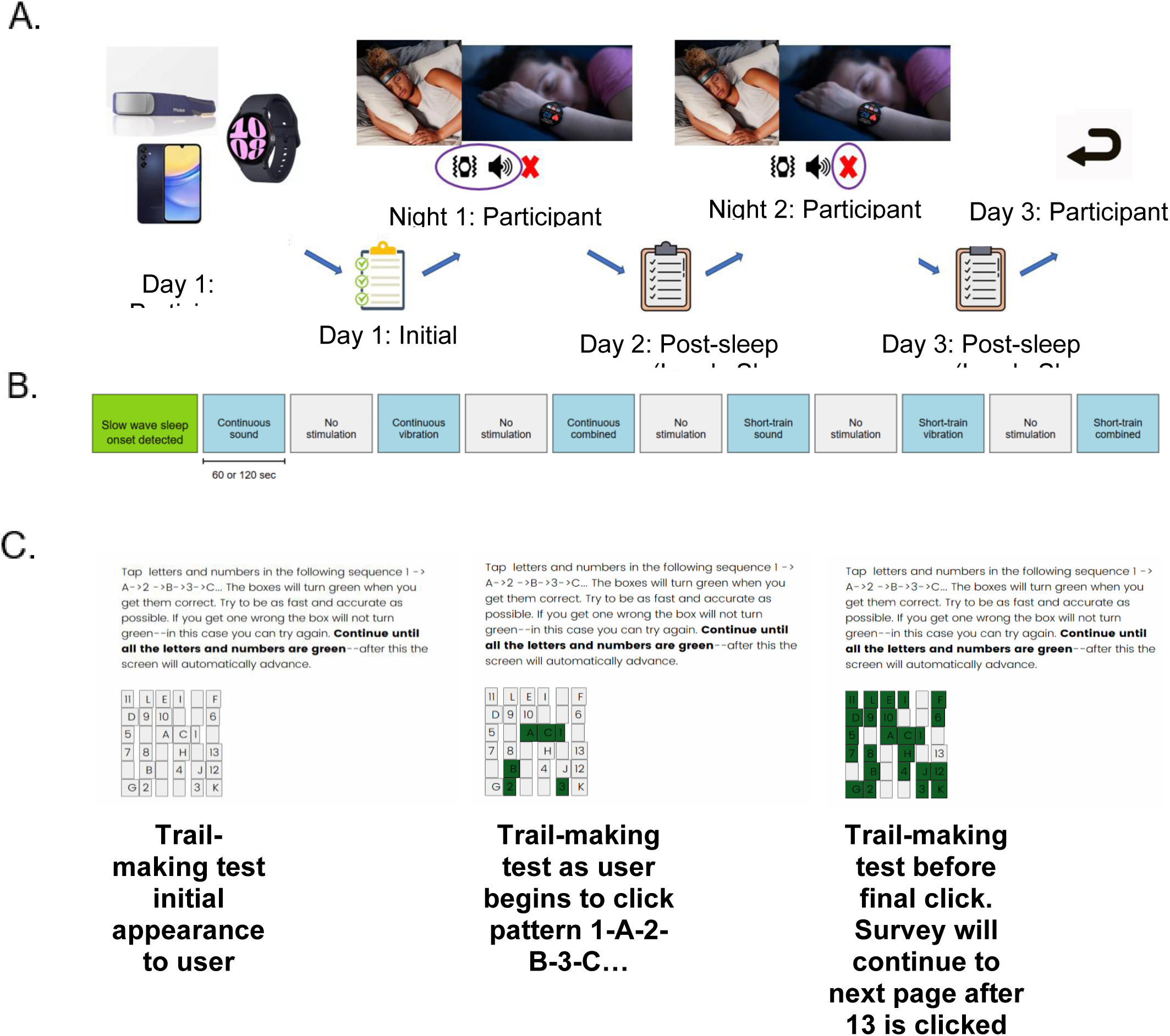
**(A)** Sequence of events in the study. **(B)** Stimulation protocol used on stimulation nights. The stimulation blocks were presented in a random order with replacement such that a participant would first receive one block of each type before repeating the blocks. The block length was 2 minutes (120 seconds) for the first group of 37 participants and shortened to 60 for the remaining 57 participants to maximize the chances of participants receiving each block type. If an arousal occurred during stimulation, stimulation was paused and restarted at the same point in the stimulation sequence once the participant again met the threshold for stimulation. **(C)** Example of how participants complete the computerized trail making task.

Therefore, there were a total of 6 stimulation conditions. Stimulation conditions were presented in blocks which lasted either 120 seconds (for the first 37 participants) or 60 seconds (for subsequent participants). After each stimulation block was an equally long no-stimulation block where no stimuli were presented. The watch continued monitoring sleep state during the stimulus and no stimulus blocks and would pause the protocol if the user left N3, resuming at the same point in the protocol once stable N3 resumed. Of the 93 participants enrolled, 68 had stimulation recorded during the stimulation night and these participants received a mean of 9 stimulation blocks. Participants completed the entire experiment at home and experimenters assisted with technical problems via text/email/ and phone. Experimenters also demonstrated how to set up the equipment when participants first visited the lab, especially focusing on how to acquire a good signal from the EEG.

When the watch detected a transition out of deep sleep (either the probability of N3 falling below 20% or a large body movement), the stimulation was immediately paused. Once the participant returned to deep sleep, stimulation was resumed in the same condition.

### Primary and secondary outcomes

Our pre-specified primary outcomes were change in mean delta event related spectral perturbation (ERSP) during stimulation (compared to same night no-stimulation blocks), difference in mean time to complete the trail making task following the stimulation night and the sham night, difference in mean click count to compute the trial making task following stimulation and sham; and change in mean LSEQ and BRUMS scores following stimulation and sham. Secondary outcome measures were the mean percentage of stimuli in N3, compared to the percentage of N3 in the sleep recording, and the mean sleep disruption rate for each stimulation type compared to non-stimulation blocks (not pre-specified).

### Randomization and blinding

Participants were randomly assigned to receive entrainment on either their first night or second night by a function in the watch app. The watch switched between stimulation and non-stimulation (sham) mode whenever the onset of a new night of sleep was detected by the Wear OS sleep detection API. Previous data, as well as our pilot testing, suggest that participants only rarely perceive stimuli targeted to deep sleep[29]; thus we compared the stimulation night to a night with no audio or tactile stimulation. Because the first night was initialized to a pseudorandom state, both participants and experimenters were blind to the order of conditions while EEG and behavioral data was collected; experimenters were unblinded to the order during analysis.

### Smartwatch-Based Sleep Detection

To deliver slow wave stimulation using the Galaxy Watch 6, we developed a custom app which ran continuously on the watches. Sound stimuli were presented using the watch’s internal speaker, and vibration stimuli using the internal vibration motor. We developed a machine learning model to predict the sleep stages given by a home PSG device (Dreem 2)[30] using the heart rate, gyroscope, and accelerometer data collected by a smartwatch. The model was trained to perform 5-stage sleep staging (Wake, N1, N2, N3 and REM) using data collected in a previous study[29] (Figure S1); the data consisted of approximately 12 million data points collected from participants ages 18-35. Receiver Operating Characteristic curve (ROC-AUC) for detecting N3 (versus non-N3) was 0.6 each second, indicating that the model detected N3 at above chance rates (Figure S2). More details about the model architecture can be found in Supplemental Note 1. When participants slept, the app estimated sleep stages once per second based on the heart rate and motion data. Stimulation began when the following criteria were met, and was paused whenever any were not met:

1. 10-minute average probability of N3 was >=25%
2. Probability of N3 in the last second was >=25%
3. No large body movements (more than 3 degree/sec motion on gyroscope reading) were detected in the last 60 seconds
4. Participant has been asleep for at least 30 minutes
5. No more than 30 min of stimulation have been delivered this night (stimulation duration was limited as little literature exists on the effects of delivering N3 stimulation for an extended period).

We applied these criteria on data held out from the original training set to characterize the stimulation stages where the model would stimulate (Figure S3). The model stimulated most often in N2 and N3 sleep and the fraction of stimulation in N3 was higher than the overall fraction of the night scored as N3, confirming the system could target N3 sleep.

### Stimulation Intensity Control

To present stimuli at effective intensity without awakening the user, we developed an adaptive algorithm for controlling the intensity of auditory and vibration stimuli. For each stimulus type, the intensity begins at 0 at the beginning of a night of sleep and increases slightly each time a stimulus is presented. If the user transitions out of deep sleep while stimuli are being presented, the system sets a cap on the intensity of that stimulus which is 95% of the intensity where the transition out of deep sleep occurred. For the rest of the night, stimulus intensity is not permitted to exceed the cap; the cap can also be decreased further if another transition out of deep sleep occurs during stimulation. Caps are set independently for auditory and vibration stimulation; for example if a user transitions out of deep sleep during vibration stimulation only the vibration intensity will be capped. Caps are reset at the end of each night.

### EEG Recording

Participants recorded their EEG while sleeping at home using an Interaxon Muse S active EEG headband linked to a smartphone provided by the lab. We analyzed data from the two frontal channels with a reference at Fpz. When participants visited the lab, we demonstrated how to use the headband and adjust it to obtain good signal quality. Participants wore the headband on both nights of the experiment. Data was recorded using the Mind Monitor app[31] and the watch was continuously synchronized to the phone’s internal clock over Bluetooth, allowing us to identify when in the EEG data stimulation occurred.

When participants visited the lab to collect equipment, we demonstrated how to put on the headband and start recording. We emphasized how to use the built in impedance monitor to obtain a stable low impedance connection on each electrode, and gave participants strategies for achieving this with the device. Participants set up the two recordings in their own homes before they went to bed.

Previous research has shown that sleep can be scored from the Muse S signals with acceptable validity[32] and spectral power from the Muse S also shows good correlation with conventional EEG[33]. However it is also well-known that dry electrode devices like the Muse have a higher noise floor at low frequencies than wet-electrode devices[34].

To address this limitation, we used an event-related analysis to examine how stimulation onsets affect delta power; which compensates for noise through baseline correction;. Event related analysis also makes it straightforward to exclude data with large artifacts.

Similarly, we chose to analyze sleep stages only in specific periods around the stimulus, rather than analyzing the entire night of sleep. We used this approach because (1) we anticipated that SWE would have an acute effect only when stimulation was on and (2) most nights contained some periods of time where sleep was not stageable, which made it impossible to give credible whole-night measures of sleep stages.

### Signal Processing and EEG Analysis

Our primary analysis examined the effects of stimulation on the delta-band EEG computed online by the Muse-S, which spans 1-4 Hz, encompassing most of the flow wave band. While the Muse delta band is not completely equivalent to the slow wave band, this band is dominated by slow wave activity around 1-1.5 Hz (due to the 1/f frequency scaling of the EEG) and therefore we considered it an adequate proxy for slow wave activity.

### Analysis of event-related delta power

For our primary analysis, we computed event-related delta power using the absolute delta power (1-4 Hz) calculated by the Muse S on channels AF7 and AF8 once per second. We used the stimulation records recorded by the watch to insert events in the Muse time series when each block of stimulation started, allowing us to compute how delta power changed when stimulation began compared to the period before stimulation. We considered the onset of each block to be one trial, and we computed the change in delta power by dividing the delta power from 0-30 seconds after the trial start by the delta power from −30 to 0 seconds before the trial start. We computed the event-related change separately for AF7 and AF8. We then computed an overall event related change for each trial by averaging the change in the AF7 and AF8 channels. Because channels can become disconnected from the head during sleep, we computed the overall event related spectral change by averaging AF7 and AF8 if both channels were connected; if one channel was disconnected then the overall value was the value of the remaining connected channel. If both channels were disconnected during a trial, that trial was excluded completely. We detected disconnected channels by looking for ERSP values of 1.0, which indicated that the channel received only noise. After obtaining a single value for each trial, we then computed the mean event related change for each participant and each stimulation condition. We then tested whether the mean ERSP for all stimulation conditions differed from the ERSP for non-stimulation condition using a paired t test. A secondary analysis performed paired t tests comparing each individual stimulus type to the non-stimulation condition. Participants were excluded from this analysis if they did not have valid EEG recordings overlapping with the delivery of stimuli.

### Sleep Stage Targeting Analysis

To determine sleep stages when stimulation was delivered, we compared the distribution of sleep stages in stimulation ON periods to the overall distribution of sleep stages. To obtain sleep stages, we bandpass filtered the raw Muse data (sampled at 256 Hz) between 0.1 and 40 Hz and re-referenced it to TP10 (approximating the standard mastoid reference used for lab based sleep staging[35]. If data were too noisy to allow for sleep staging, data were referenced to a different electrode (either TP9, AF7, or AF8) and/or additionally filtered with a lowpass filter at 25 Hz. Data were then epoched from −60 to +30 seconds relative to block starts. For our primary analysis, a trained sleep scorer determined the AASM sleep stage[35] in the −30-0 epoch relative to the beginning of stimulation blocks. This time period was chosen because it represents the sleep stages in which the system initiated stimulation, uninfluenced by changes in sleep induced by stimulation. To determine if our system successfully targeted N3, we also constructed a baseline distribution for the prevalence of each sleep stage by randomly sampling 10 90-second windows from 17 night recordings with stagable Muse EEG. We used the middle 30 seconds of each window to construct a randomly baseline distribution of sleep stages for our participants. Only the middle 30 seconds were used as accurate sleep staging requires also staging 30 seconds before the epoch to be analyzed and 30 seconds after. We then used a chi squared test to test whether the percentage of stimulation in each sleep stage differed significantly from the proportion of that stage in the baseline. A significant chi-square would indicate that the system either targeted in that stage or avoided stimulating in that stage. Participants were excluded from this analysis if they did not have EEG judged as manually stagable overlapping with stimulation.

### Sleep Disruption Rate Analysis

To measure whether stimulation disrupted sleep, we measured the “disruption rate” for each stimulation condition and the no-stimulation condition. While conventionally, sleep disruptions would be defined by looking for shifts in EEG activity[36], we found the home EEG signal was poorly suited for this analysis due to low signal to noise ratios and artifacts related to movement when sleep is disrupted. We defined the disruption rate using smartwatch data as the number of times the participant transitioned from an “N3 detected” state to an “N3 not detected” state per block, for each block type. This metric therefore provides a measurement of sleep disruptions per unit of time, similar to common EEG-based metrics like the sleep fragmentation index[37]. We compared disruption rates between block types using a paired t test within each participant. Participants were excluded from this analysis if they did not have stimulation recorded on their stimulation night.

### Behavioral and Survey Measures

To examine the psychological and cognitive effects of slow wave stimulation, we collected several surveys after each night of sleep and compared results after the sham night to results after the stimulation night. All surveys were emailed to participants 24 and 48 hours after they visited the lab and completed at home on a computer or phone.

Sleep quality was obtained using the Leeds Sleep Evaluation Questionnaire (LSEQ)[26], which asks participants to rate the ease of falling asleep, quality of sleep, ease of waking up, and feeling during the day. We measured all subscales of this questionnaire except for the Getting To Sleep scale, which we considered not relevant since stimulation began after sleep onset. Participants also completed the Brunel Mood Scale (BRUMS)[27], which evaluates their current state on 7 axes (tension, depression, anger, vigor, fatigue, and confusion), plus a composite axis (overall negative mood) which reflects the sum of all other axes minus vigor.

To evaluate cognitive function, we developed an online version of the Trail Making Test part B (Trails B), a test of planning, psychomotor speed, and executive function that is sensitive to sleep deprivation [38,39]. Trails B requires participants to draw a line connecting a series of randomly spaced letters and numbers in the order 1 -> A -> 2 -> B…etc. The primary measure obtained from this task is the time taken to complete it.

Trails B is typically a pen and paper task; to adapt it to an online study we created a computerized version of it (Figure 1C). In our modified trails B, 13 letters and 12 numbers are randomly distributed on a 6 x 6 grid. Participants are asked to begin with clicking the 1 button, and then A, continuing to 2->B until reaching L. Each letter/number turns green if it is clicked in the correct order; if a participant clicked an incorrect letter/number they must find the correct one and tap it to continue. We record the total time to complete the task, and also the number of clicks required to complete it.

### Evaluating the sleep quality, mood, and cognitive effects of slow wave entrainment

To measure whether slow wave entrainment affected sleep quality, mood during the day, or performance on the Trails B task, we compared test performance after stimulation and no-stimulation nights using a paired t test. Participants were excluded from these analyses if they did not have one night of stimulation and behavioral measures following both the stimulation night and sham night.

An ancillary pre-specified analysis tested whether specific stimulation parameters and individual factors predicted the effects of stimulation. For these analyses, we used linear regression to test whether each factor predicted the difference in test score following the stimulation night and test score following the sham night. Each combination of factor/test was tested individually (i.e., we did not perform multiple regression) as we expected factors like age, mental health,and baseline sleep quality to be highly collinear.

## Results

### Slow wave entrainment increased EEG delta power

We found that frontal delta power (1-4 Hz in electrodes AF7 and AF8) increased when slow wave stimulation was initiated compared to a 30-second baseline before each stimulation period [one sample t test; t(34)=3.04; P =.005]. Delta power also increased during stimulation blocks (Stimulation on) compared to the non-stimulation (Stimulation off) blocks [paired t test, t(32)=2.55, P = .02] (Figure 2A).

**Figure 2:**
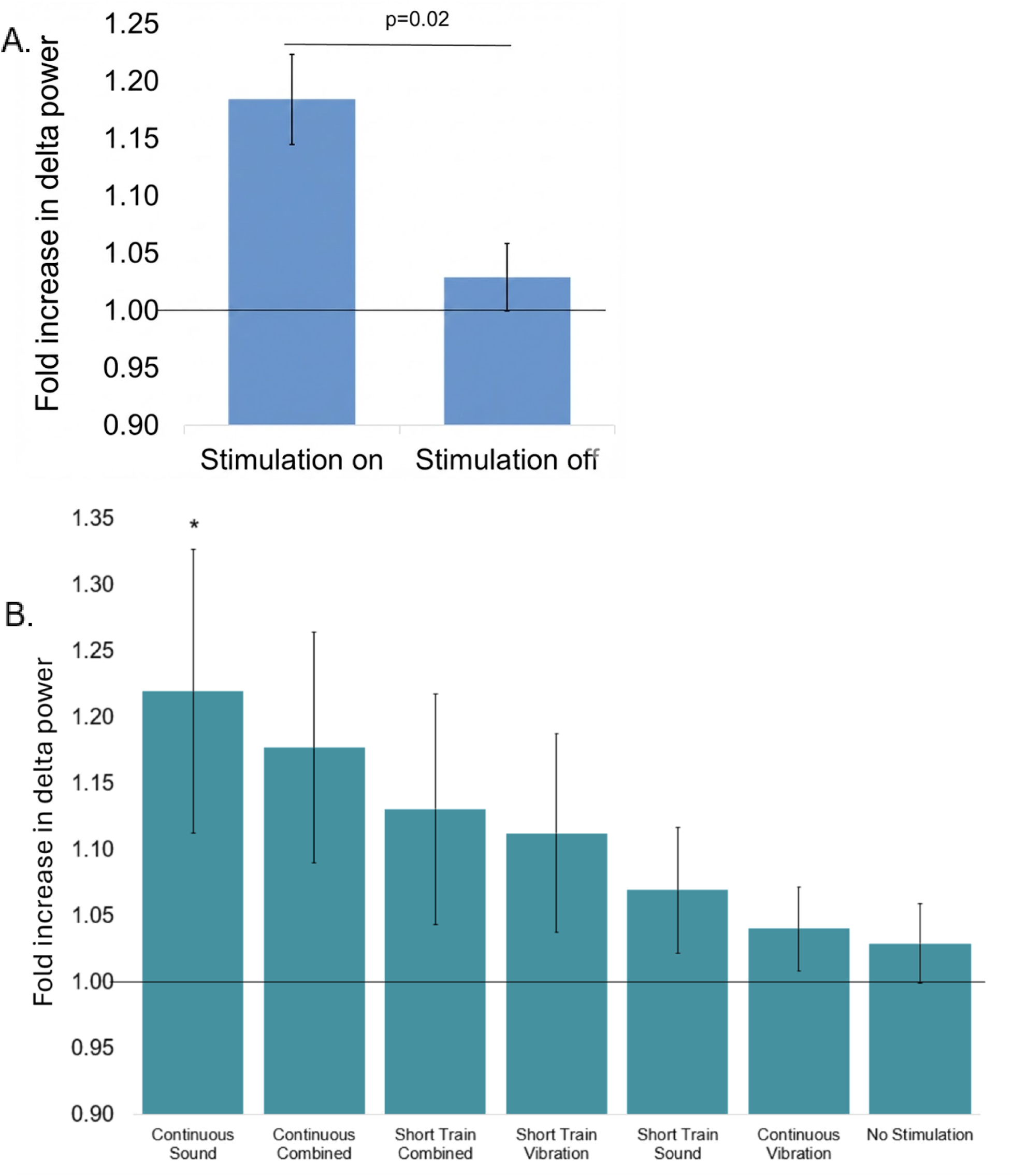
**(A)** Effects of slow wave stimulation on frontal (AF7/AF8) delta power quantified using event related spectral perturbation (ERSP), quantified as delta power during block/delta power over the previous 30 seconds. Delta power evoked by the stimulation was significantly greater than delta power evoked by the non-stimulation block. Both the stimulation and non-stimulation blocks had a significant increase in delta power over baseline (indicated by the horizontal line at Delta ERSP = 1.00); in the non-stimulation block this likely resulted from the natural increase in delta power as participants progressed into deeper N3 sleep, as delta amplitude slowly increases during transitions into N3[40] Data are for 35 participants with delta band data available during stimulation. **(B)** Effects of specific stimulation types on frontal delta power. Asterisks indicate a significant power increase (p < .05) over baseline.

We also tested the effects of different stimulation modalities (sound, vibration and combined sound/vibration) and duty cycles (continuous and short train) in inducing delta waves (Figure 2B). The effects of different stimulus modalities did not differ significantly, with all showing an increase in delta power over baseline.

### Slow wave entrainment decreased sleep disruptions

Presenting stimuli in sleep carries a risk of increased sleep fragmentation, which can decrease the overall quality of sleep[41]. To determine whether stimulation increased sleep disruption, we measured the proportion of blocks that contained a smartwatch-detected transition out of stimulatable sleep for each condition, which we called the disruption rate. We found that disruption rate was significantly lower for stimulation blocks than for non-stimulation blocks (Figure 3).

**Figure 3:**
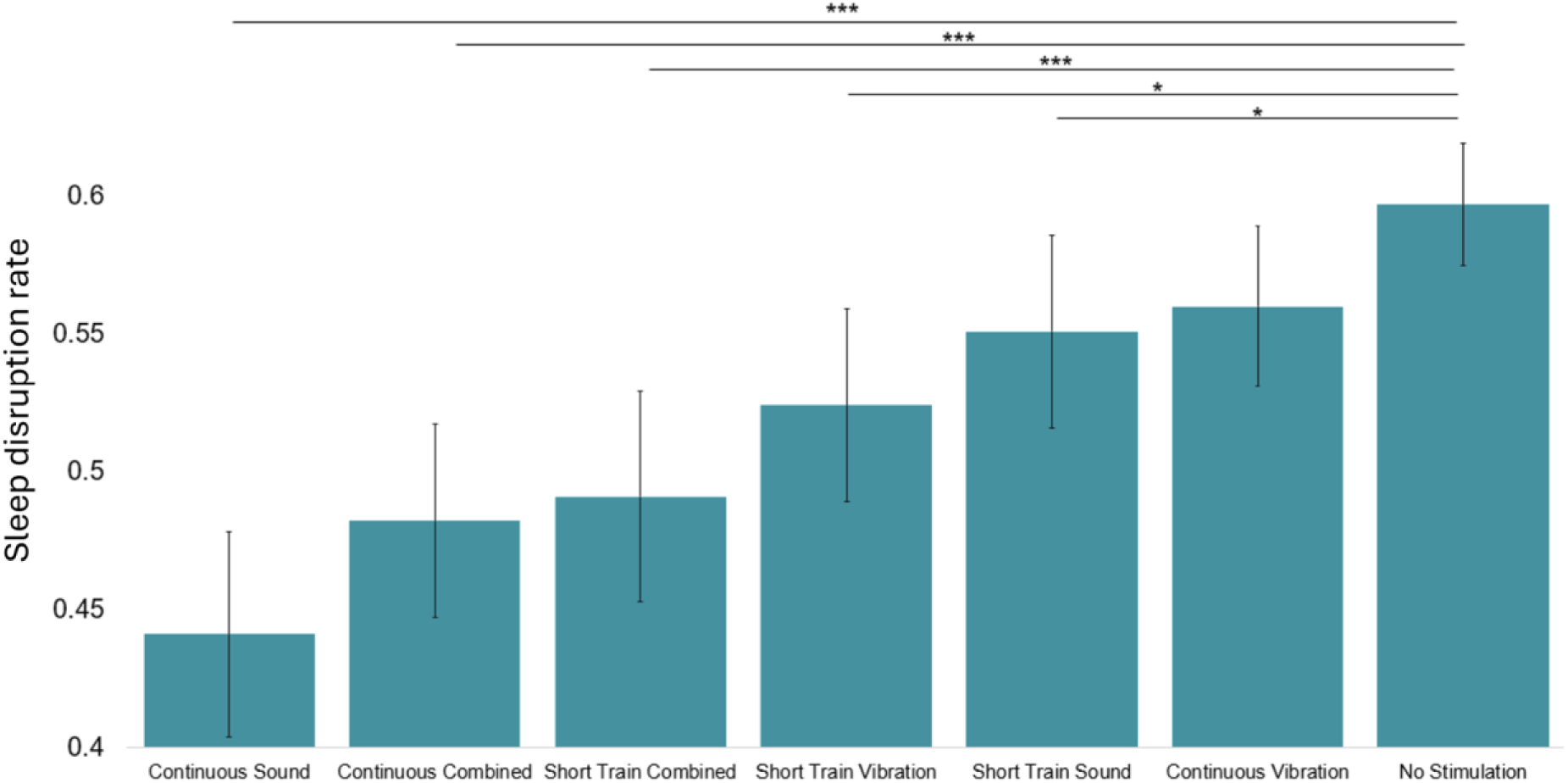
The effect of stimulation condition on sleep disruption rate. Asterisks represent significant differences in arousal rate using a two-tailed t test (*: p < .05, ***: p < .001). All stimulation conditions other than the continuous vibration condition had a significantly lower arousal rate than the no-stimulation condition. Data are for 68 participants who had stimulation data recorded.

### Slow wave entrainment improved cognitive function and sleep in participants with large delta ERSPs

We did not find any significant main effects of stimulation on our measures of sleep quality, mood, and cognition. However, the degree of delta wave enhancement that occurred with stimulation predicted cognition and sleep improvements in the stimulation night; in particular, the size of the frontal delta ERSP to stimulation was associated with improvements in sleep quality, mood, and Trail Making Task performance (Table 1, Figure 4) in a two-tailed linear regression.

**Table 1:**
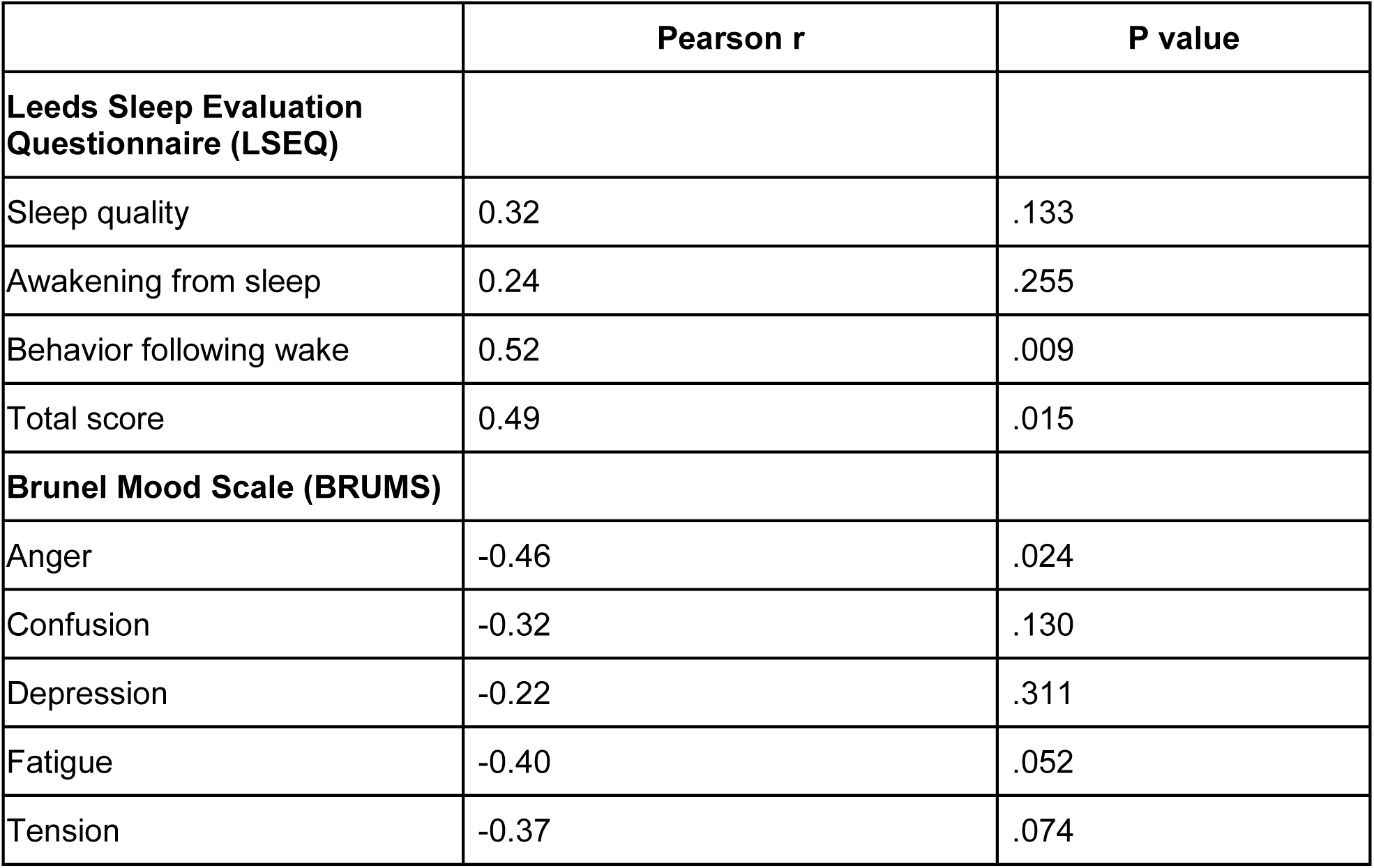
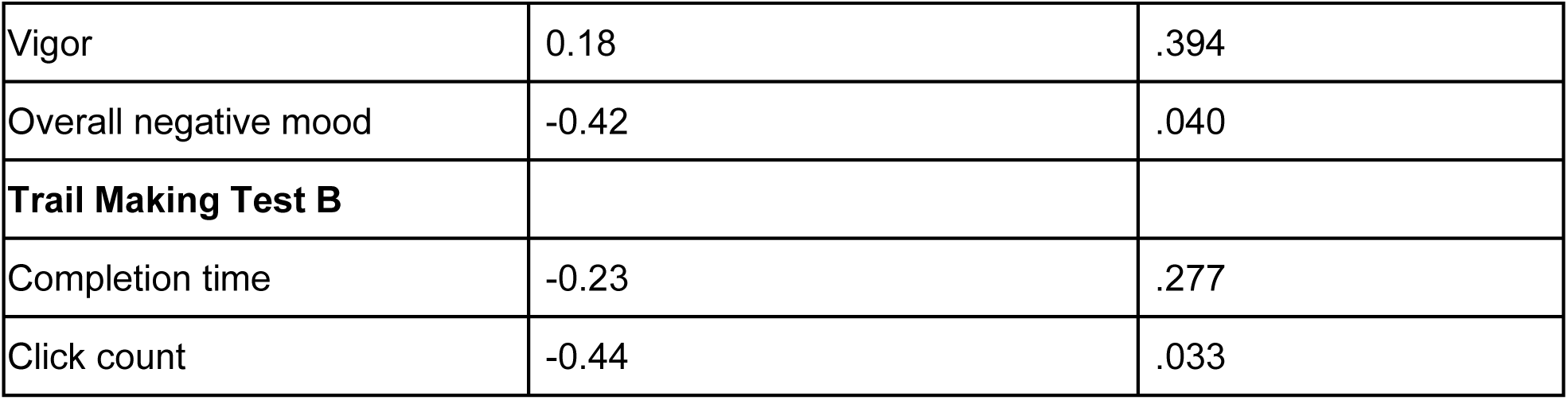
Correlation between the change in behavioral outcome measures from stimulation night to sham night and the increase in delta power evoked by stimulation on the sham night. A positive r value indicates that the stim-sham value was positively correlated with the amount of delta power induced by the stimulation. We found that increased delta power was associated with better self-reported sleep quality (LSEQ behavior following waking, total LSEQ), reduced BRUMS anger, and reduced overall negative mood on the BRUMS). We also found increased delta power predicted a reduced number of clicks needed to complete the Trail Making Task, indicating a reduction in the number of errors on this task (since the Trail Making Task requires a minimum of 26 clicks to complete it successfully). Data are for 24 participants who had valid EEG on both nights, stimulation on their stimulation night, and completed questionnaires after both nights.

**Figure 4:**
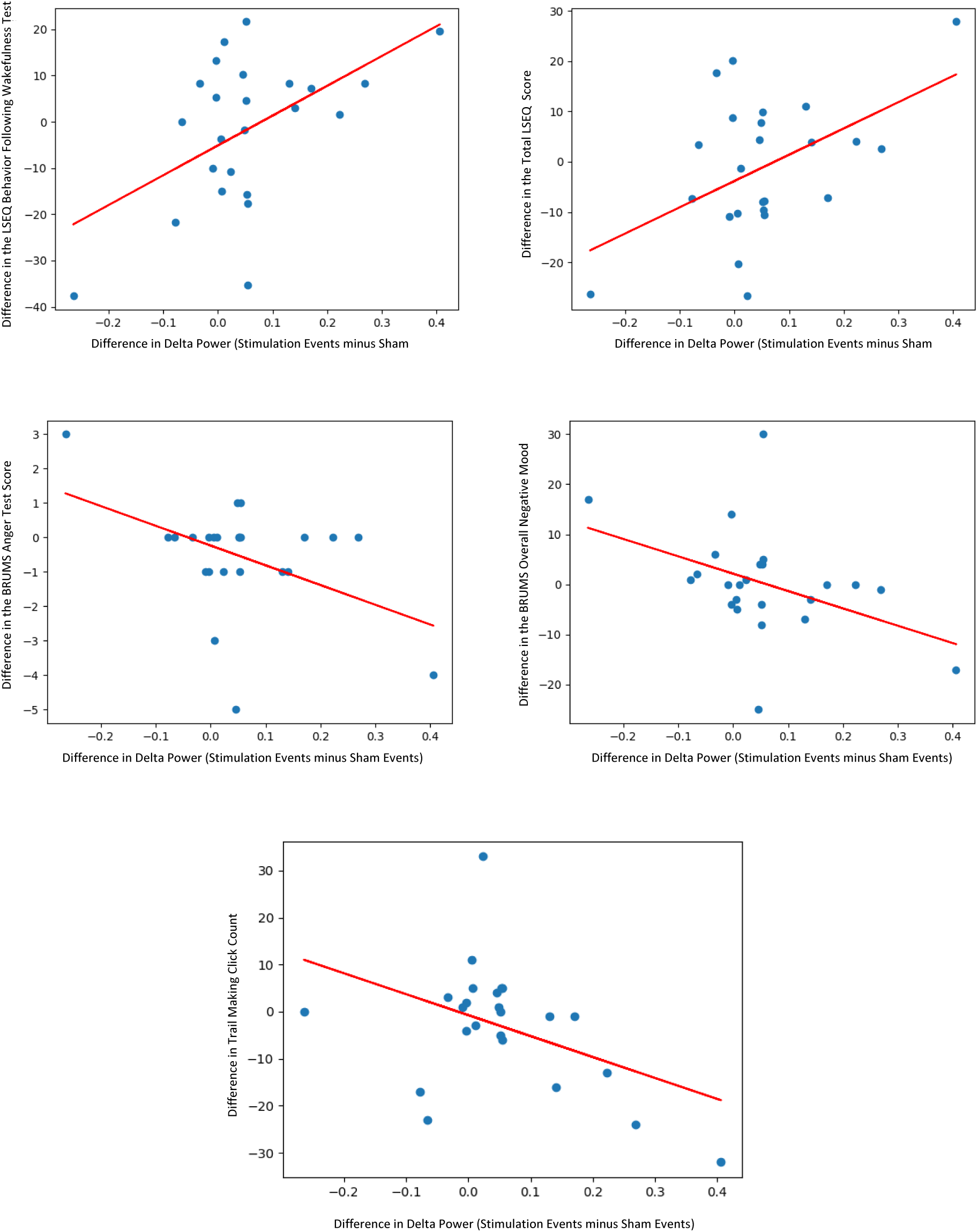
Scatter plots of the significant correlations in Table 1. The difference on the Y axis reflects score after the stimulation night minus score after the no-stimulation night.

### Slow wave entrainment improved mood in participants with poor baseline mental health

We tested whether the difference between the stimulation night and non-stimulation night was correlated with participant factors including age, typical bedtime, typical sleep duration,Pittsburgh Sleep Quality Index (PSQI) PSQI and General Health Questionnaire (GHQ-12) scores. We found that participant baseline mental health (GHQ-12 score) modulated the effects of stimulation on confusion and tension mood scales; participants who reported more mental health problems on the GHQ-12 at the start of the experiment showed a larger reduction of tension and confusion after their stimulation night (Table 2, Figure 6). Participant age, typical sleep duration, typical bedtime, and PSQI score did not significantly modulate the effects of slow wave entrainment.

**Table 2:** Correlations between participant demographics and the effects of slow wave stimulation. Effects of the stimulus on all measures were computed by calculating the difference (stimulation night - no stimulation night) for the metrics and are based on 41 participants who completed the initial questionnaires and both nightly questionnaires.

| Outcome measure | Age r (P) | Bedtime r (P) | Sleep duration r (P) | GHQ-12 r (P) | PSQI r (P) |
| --- | --- | --- | --- | --- | --- |
| **Leeds Sleep Evaluation Questionnaire (LSEQ)** |  |  |  |  |  |
| Sleep quality | 0.01 (.94) | 0.20 (.22) | 0.06 (.71) | -0.04 (.79) | 0.02 (.90) |
| Awakening from sleep | 0.03 (.88) | 0.03 (.83) | 0.04 (.81) | -0.00 (.98) | 0.00 (1.00) |
| Behavior following wake | 0.10 (.54) | 0.08 (.61) | 0.04 (.82) | 0.03 (.85) | 0.03 (.86) |
| Total score | 0.05 (.74) | 0.16 (.33) | 0.06 (.69) | -0.02 (.92) | 0.02 (.89) |
| **Brunel Mood Scale (BRUMS)** |  |  |  |  |  |
| Anger | -0.23 (.16) | 0.02 (.90) | 0.21 (.18) | -0.28 (.08) | -0.18 (.25) |
| Confusion | -0.22 (.18) | 0.08 (.60) | 0.18 (.25) | -0.34 (.03) | 0.04 (.83) |
| Depression | -0.22 (.18) | -0.01 (.95) | 0.24 (.12) | -0.18 (.25) | -0.00 (.98) |
| Fatigue | -0.11 (.51) | -0.11 (.49) | 0.08 (.61) | -0.02 (.93) | 0.01 (.95) |
| Tension | -0.19 (.22) | 0.18 (.25) | 0.20 (.20) | -0.46 (.003) | -0.02 (.92) |
| Vigor | 0.04 (.82) | 0.22 (.18) | 0.03 (.84) | -0.04 (.81) | -0.13 (.42) |
| Overall negative mood | -0.21 (.20) | -0.06 (.69) | 0.17 (.28) | -0.22 (.18) | 0.01 (.93) |
| **Trail Making Test B** |  |  |  |  |  |
| Completion time | -0.07 (.66) | -0.07 (.67) | 0.00 (.98) | 0.09 (.57) | 0.06 (.71) |
| Click count | -0.02 (.92) | -0.12 (.44) | -0.04 (.79) | 0.12 (.45) | 0.07 (.66) |

**Figure 5:**
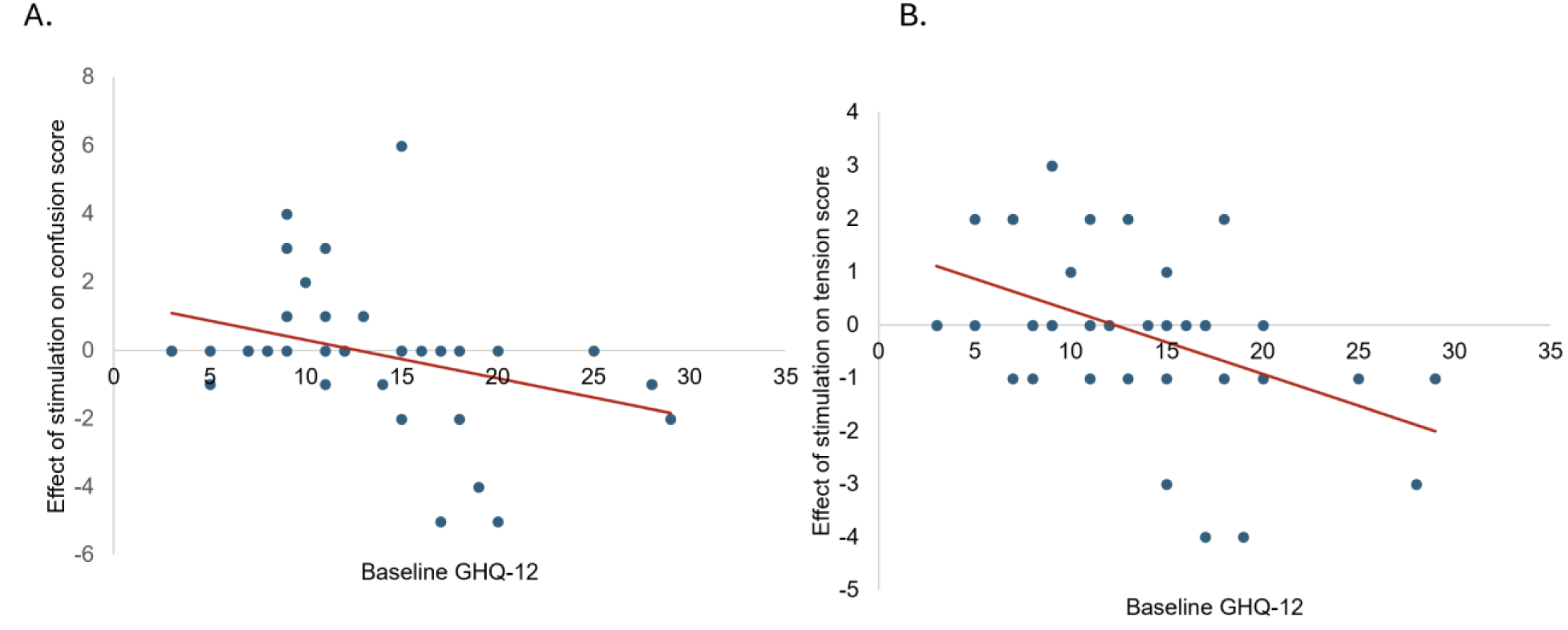
The significant correlations between initial GHQ-12 score and effects of the stimulation on BRUMS confusion and tension axes. The Y axis represents the value on the stimulation night minus the value on the sham night.

### Entrainment stimuli targeted N3 sleep

To assess which sleep stages the system targeted for stimulation, a trained sleep scorer performed manual sleep staging on the −30 to 0 second period relative to the start of stimulation blocks using the AASM criteria[35] and also staged 10 randomly selected 90 second periods of sleep in each night to create a baseline distribution of sleep stages. We considered only the period before stimulation started in order to measure which phases of sleep the system targeted for stimulation without the confound of changes in sleep physiology induced by the stimulation. We obtained baseline sleep stage distributions for 17 nights which had stageable Muse EEG and determined sleep stages of stimulation onsets in 10 nights which had both Muse EEG data and stimulations (Figure 6). Stimulation was significantly more likely to occur in N3 compared to the baseline distribution (odds ratio=1.75, X^2^(1, 229)=5.01, P=.025) and was significantly less likely to occur in wake (odds ratio=0.18, X^2^(1, 229)=10.86, P=.001) as shown in Figure 7. The rate of stimulation in N1, N2, and REM sleep did not differ significantly from the baseline occurrence of those stages. Despite targeting N3, stimulation occurred most frequently in stage N2 (33% of stimulus onsets) due to the higher base rate of N2 compared to N3. Stimulation occurred next most frequently in N3 (29% of onsets) REM (28% of onsets), N1 (8% of onsets) and wake (3% of onsets).

**Figure 6:**
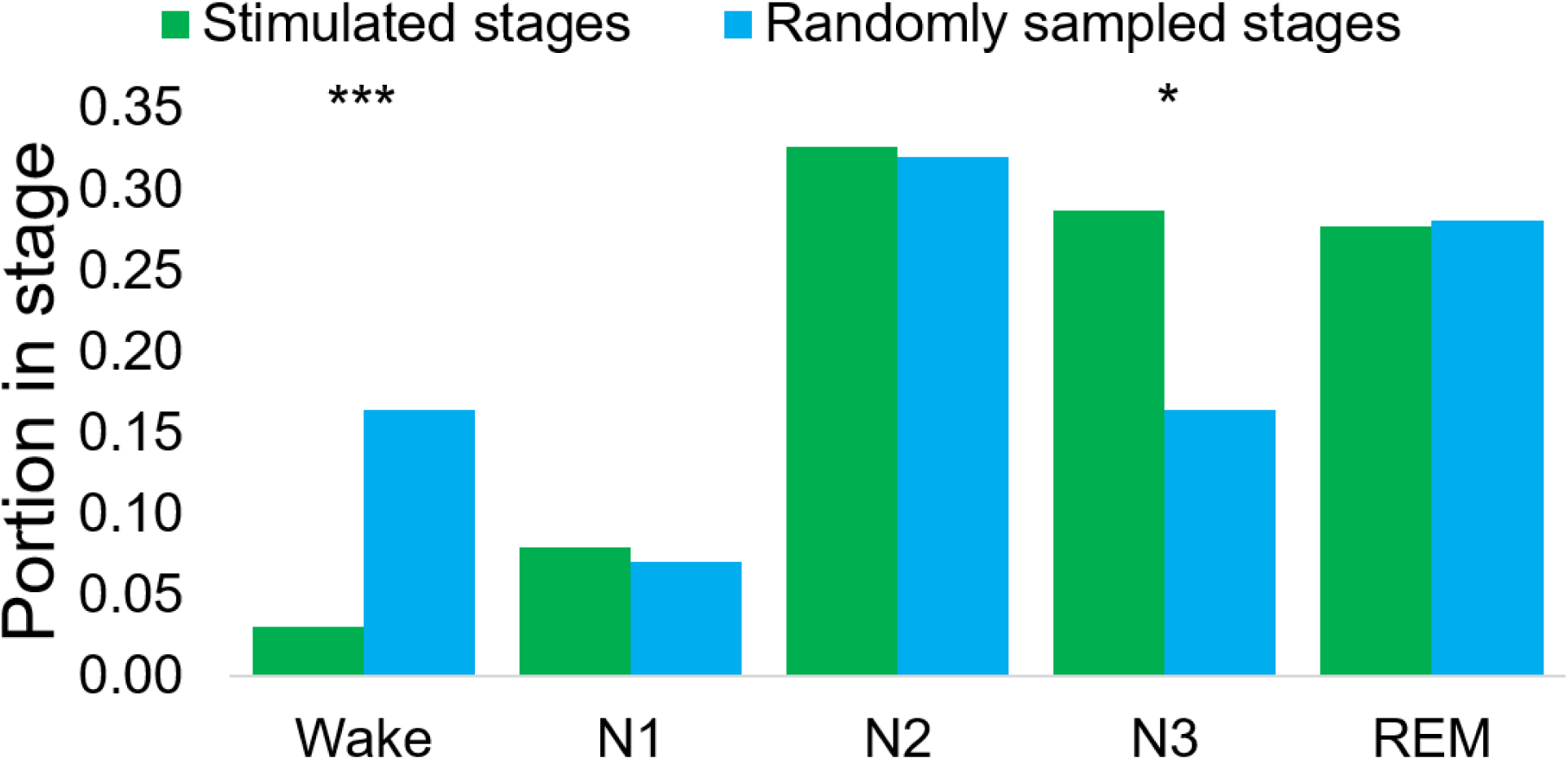
Portion of the stimulation delivered in each sleep stage in our manually staged subset. Asterisks indicate significant differences (chi-square p < .05) where stages were overrepresented or underrepresented in the stimulated periods compared to the overall distribution of stages. Data for the randomly sampled stages are from 17 nights that had stagable EEG data and data for the “stimulated stages” bars are from 10 nights which had both stagable EEG data and stimulations.

### Participants rarely reported perceiving entrainment stimuli

Out of 86 participants who completed the post-sleep questionnaire, 12 reported perceiving sound or vibration stimuli at some point during the night and of these 5 reported that the stimuli disturbed their sleep. The remaining participants who perceived stimuli reported that they occurred either while falling asleep or when they had awakened for some other reason.

A participant flow diagram is shown in Figure 7.

**Figure 7:**
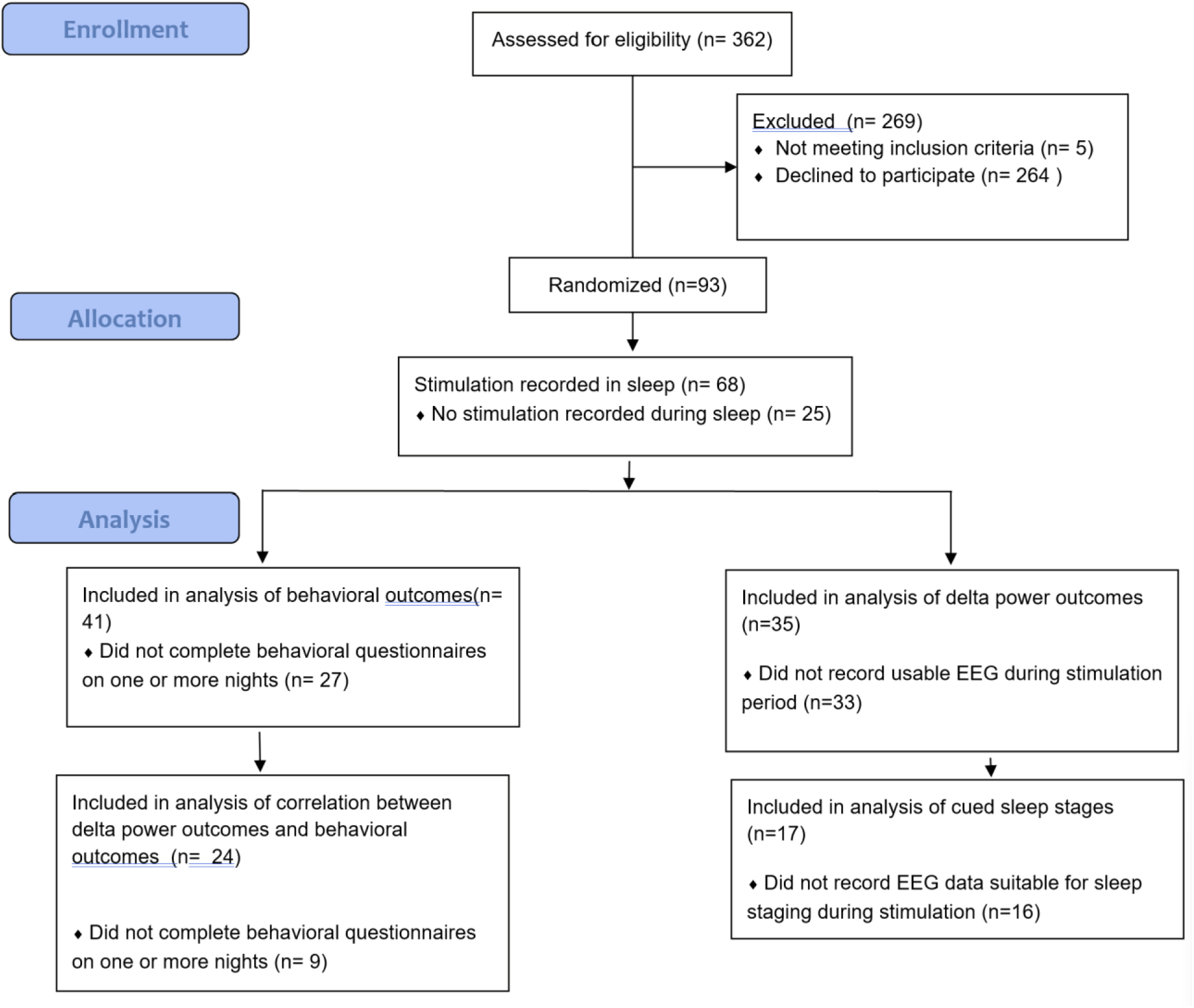
CONSORT diagram of participant flow through the experiment. Because we used a within-subjects manipulation, participants were not allocated to groups.

## Discussion

### Principal Results

In this study, we found that rhythmic auditory and vibrotactile stimulation delivered by a smartwatch was associated with increases in frontal delta power during sleep.

Although we did not observe significant group-level differences between stimulation and no-stimulation nights on measures of sleep quality, mood, or cognitive performance, we found that the degree of delta wave enhancement that occurred with stimulation predicted cognitive and sleep improvements across individuals. Stimulation was associated with increases in delta power across multiple stimulus and stimulation patterns, and we did not observe significant differences between the different stimulus modalities, implying that all had similar effects.

While we did not observe overall differences in cognition and subjective sleep quality after stimulation nights compared to control nights; we did observe significant correlations between the behavioral effects of stimulation and the amount of delta power elicited by stimulation. This finding suggests that people varied substantially in how well they responded to entrainment, but participants who showed a strong neural response to entrainment also showed a strong behavioral response to entrainment. Notably, this correlation cannot be explained by a correlation between trait delta power and trait sleep quality, because we found that the *difference* in delta power when stimulation turned on explained the difference in outcomes between two counterbalanced nights. Our results thus suggest that the behavioral benefits of slow wave entrainment depend on increasing delta power, and when this is successful participants may experience improvements in mood, sleep quality, and cognition.

Several factors may explain why participants vary in how well they respond to stimulation. The ability of the brain to entrain to sensory stimuli may vary with demographic factors like age[42,43]. Additionally, participants who are highly arousable may experience sleep disruption rather than benefit from the stimuli, as evidenced by our findings that some participants with low evoked delta showed a decrease in behavioral measures after stimulation. Finally, a participant’s baseline state may dictate how much improvement is possible, as shown by our findings that stimulation was more effective at improving mood in participants with poor baseline mental health.

We also found evidence of improved sleep stability stimulation relative to matched no-stimulation periods. Slow-wave entrainment may reduce arousability during sleep, as slow-wave sleep and sensory-elicited K complexes have both been linked to deep, unarousable sleep, and K complexes are suspected to play an active role in suppressing sensory stimuli that would otherwise cause awakening[44,45].

### Comparison With Prior Work

A unique feature of this work is that SWE was delivered using a smartwatch which sensed sleep to determine the optimal timing and intensity of stimuli. Compared to previous EEG-based methods, using a smartwatch can improve the accessibility of SWE, while also retaining the ability to monitor biosignals and avoid disrupting sleep.

Because smartwatches cannot detect the phase of individual slow waves, stimulation was delivered without phase locking to individual slow waves. Most prior SWE studies have employed phase-locked stimulation where sounds are locked to an optimal phase of the ongoing slow oscillation[18] though previous studies have also shown that non-phase locked SWE can improve sleep parameters[23,46]. Notably, no studies have combined non-phase-locked entrainment with an algorithm that targets periods of deep sleep and modulates the stimulus intensity to avoid disturbing sleep. Our findings support the idea that SWE which is not phase locked but is targeted to deep non-REM sleep can increase delta oscillations and produce beneficial effects on sleep.

Prior research has found that slow-wave entrainment may be less effective in older adults than in younger adults[42,43,47]. By contrast, our study found that age did not modulate the degree of entrainment or the effects of slow wave entrainment on mood, cognition, and sleep quality, despite including participants across ages 18-92. One possibility is that entrainment with non-phase-locked stimuli may be less affected by age because it does not depend on accurately detecting the peaks of slow waves, which are less prominent in older adults.

Manual sleep staging based on EEG indicated that the proportion of stimuli presented in N3 was higher than expected by chance (Figure E4), although stimulation also frequently occurred in N2, and, less commonly, REM and wake. This is likely because we used a liberal N3 detection threshold in this experiment to ensure that participants received sufficient stimulation to enable comparisons across experimental conditions; increasing the threshold would increase N3 specificity.

Despite stimulating outside N3 (and sometimes in wake) we found no evidence of sleep disruption by stimuli and very low rates of participants reporting noticing the stimuli. This can likely be explained by two factors: The algorithm calibrates stimulus intensity to the user’s arousability, and stops stimulation when sleep disruption is anticipated; thus it may have been able to avoid disrupting sleep even when stimuli were presented in off-target sleep stages. Secondly, SWE in stages with low natural slow wave power like N1 or REM may have a neutral or even beneficial effect [48] so long as it is not disruptive.

### Limitations

Several limitations should be considered when interpreting these findings. Because we collected data in participants homes, there were high rates of incomplete data resulting from EEG signal quality and participant adherence issues.

Participant loss resulted from several sources. 25 participants did not have any stimulation recorded due to a number of reasons including withdrawing from the study before the stimulation night, failure to properly charge and wear the watch, and no sleep that met the conditions for stimulation on the stimulation night. We had further attrition when considering the participants who had both stimulation and EEG data available (35/93); loss here was mostly attributable to challenges with EEG setup and EEG signal quality. Finally, we saw attrition due to occasional failure to complete the surveys after a night of sleep.

In this study we measured delta power changes locked to stimulus onset, but did not compare delta power throughout the night or assess the changes in sleep stage distribution throughout the entire night. This was a deliberate decision; as we hypothesized that the effects of slow wave stimulation were acute and occurred during the small fraction of the night (around 12.6 min on average) where stimulation was active. Future experiments should explore the effects on whole night sleep physiology with larger samples and increased stimulation durations.

Additionally, while we found that the stimulation targeted N3 at rates significantly above chance, there were also significant numbers of cues in other sleep stages, including light sleep and wake, which diverges from typical SWE protocols. Despite the presence of non-N3 stimulation, we observed reliable increases in delta- and slow-wave amplitudes. Besides N3, slow waves are also found in N2 and REM sleep[49,50], and it is possible our stimulation benefited sleep by entraining slow waves in these stages. We also found that the stimuli were rarely perceived (despite some stimuli occurring in wake or light sleep), likely because when stimuli were delivered outside of deep sleep the control algorithm quickly stopped estimation. Our results suggest that slow wave entrainment may be effective even when not completely selective for N3, so long as sleep disruption can be limited.

Finally, while we found interesting correlations between the degree of stimulation-induced power increase and the behavioral effects of stimulation, this analysis was exploratory, not corrected for multiple comparisons, and did not find overall behavioral effects of stimulation. Future studies should examine the cognitive, sleep quality, and mood effects of smartwatch-SWE in larger samples.

## Conclusions

This study demonstrates the feasibility of using smartwatch-based sensory stimulation to elicit slow-frequency EEG responses during home sleep. Wearable-guided stimulation paradigms may enable new experimental approaches for studying sleep-related neural dynamics outside laboratory settings[51]. While the present findings do not establish behavioral or clinical efficacy, they motivate further large scale studies into how slow wave entrainment on a smartwatch can be used to improve sleep quality.

## Code availability

An implementation of our stimulation software is available at [52]

## Data availability

De-identified raw data and scripts to recreate the analyses are available at [53]

## Acknowledgements

We are thankful for the time and trust of our participants and Stanley Buchin’s work in coordinating research at the Lasalle retirement community. This study was funded by a grant from Samsung Electronics (“New Digital Biomarkers and Interventions to Monitor and Improve Sleep and Mental Wellbeing”).

## Abbreviations

SWE: slow wave entrainment
NREM: Non-rapid eye movement
ERSP: event-related spectral perturbation
PSQI: Pittsburgh Sleep Quality Index
LSEQ: Leeds Sleep Evaluation Questionnaire
BRUMS: Brunel Mood Scale

