## Supplemental material for "Slow Wave Entrainment Using a Smartwatch: A Randomized Crossover Study"

**Supplementary material**

**Supplementary Table 1: Demographics of subgroups included in analyses**

| Group | Percent male | Mean age | Median age | SEM age |
| --- | --- | --- | --- | --- |
| Participants with delta power data | 30.30 | 43.7 | 38 | 4.24 |
| Participants with slow wave data | 25.93 | 39.93 | 36 | 4.13 |
| Participants with arousal rate data | 33.33 | 44.15 | 38 | 2.97 |
| Participants with correlations between questionnaires and demographics | 26.83 | 38 | 37 | 2.71 |
| Participants with correlations between questionnaires and delta power | 25.00 | 36 | 32 | 3.99 |

**Supplemental Note 1: Description of the real time sleep intervention system**

*Sleep staging model*
The sleep staging algorithm (summarized in Figure S1) processes data from wearable sensors to classify each second into one of five stages: wake (S0), S1, S2, S3, and REM. It utilizes three types of sensor data: heart rate, accelerometer, and gyroscope readings (each with X, Y, and Z axes). The algorithm's first step is to transform this time-domain sensor data into the frequency domain using Fast Fourier Transforms (FFTs). An FFT is calculated for the heart rate, and also for the accelerometer signal summed across all axes and the gyroscope signal summed across all axes. These FFTs are calculated over 64-sample windows (approximately one minute of data) at a 1 Hz frequency, resulting in 64 features each for heart rate, accelerometer (sum of X, Y, and Z axes), and gyroscope data (sum of X, Y, and Z axes). In addition, the values of each accelerometer and gyroscope axis in the previous second are also given as input, for a total of 199 features.

This frequency-domain representation serves as input to a neural network classifier. The network consists of four layers: three dense hidden layers with 128, 64, and 32 units respectively, each using ReLU activation functions, followed by a dense output layer with 5 units and a softmax activation. This structure allows the model to learn complex, non-linear relationships between the frequency-domain features and sleep stages.

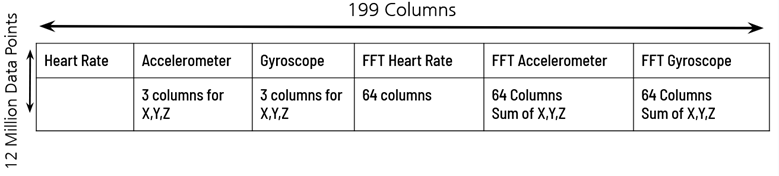

*Figure S1: summary of the sleep staging model*

We trained the model using a dataset obtained from a previous study^46^. The full dataset consisted of ~12 million datapoints sampled once per second, acquired using the procedures described in^46^. Briefly, gyroscope, accelerometer, and heart rate data were acquired from a Fitbit smartwatch while participants slept in their homes and received targeted memory reactivation controlled by the Fitbit. The Fitbit wearable data were co-registered with sleep stages obtained from a Dreem 2 home polysomnography device^52^. This process thus allowed us to train the model to predict the sleep stage from the motion (gyroscope+accelerometer) and heart rate data.

We evaluated the model by holding out the last 20% of the data from training and using it as a testing set. This allowed us to evaluate the model’s predictive abilities on participants never seen before, as well as previously seen participants. Model performance was quantified using confusion matrices representing correct and error rates for each second of sleep, as shown in Figure S2.

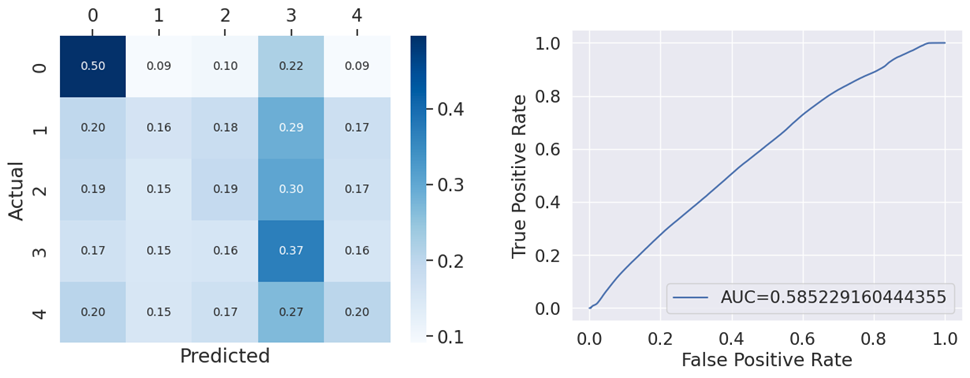

*Figure S2:*  Performance of the sleep staging model in staging each second of sleep. (Left) Confusion matrix comparing the outputs of the model against the true sleep stages determined by the Dreem 2. Stage 0 is wake, stage 4 is REM, and stages 1-3 represent N1-N3. (Right) A ROC curve of the machine learning model in predict stage N3.

*Post-processor*

While the model output can be used directly (the probability of each sleep stage is saved every second and can be used to collect statistics on sleep), it can also be used to control stimulation in sleep.

With stimulation, the goal is generally to present stimuli in a particular sleep stage. The targeted sleep stage is stage N3 (otherwise known as stage 3 sleep). The post-processor thus allows stimulation when several criteria are met:

 1. The probability of stage 3 in the current second is above a specified threshold

2. The average probability of stage 3 over the past X seconds is above a specified threshold (both values can be set by the user)

3. It has been at least X seconds since sleep onset (sleep onset is detected by the Wear OS Health Services platform)

4. It has been at least X seconds since the last detected arousal

5. No more than X stimuli have been presented

6. The user has been in a stable sleep state (defined using the criteria above) for at least X seconds

7. No more than X seconds have elapsed since the start of sleep

Arousals are defined when a large motion is detected on the gyroscope sensors, or the probability of stage 3 in the current seconds drops below its threshold, or the average probability of N3 drops below its threshold.

The post-processor controls the start and stop of stimulation, and also the stimulation intensity. Intensity is set using an automated calibration algorithm; it starts at 0 and increases linearly whenever stimulation is turned on. If an arousal is detected while stimulation is on, the intensity of the stimulation is capped at the value that triggered the arousal, minus a small offset. Stimulation then continues at this intensity until either the end of the sleep epoch (where the cap is reset) or until another arousal occurs (at which point the cap and intensity are decreased further).

*Offline system evaluation*

We performed an additional evaluation testing whether an entire system consisting of the machine learning model and the post-processor tuned to detect stage 3 sleep would accurately target stimulation to stage 3. This analysis supplements our previous analysis which evaluated the machine learning model’s ability to stage each second of sleep by demonstrating in which sleep stages would stimulation occur, taking into account both the performance of the model and the performance of the sleep staging system.

We performed this evaluation on the same validation set used for the testing of the model. Because the post-processor was optimized to target N3, we tested whether the combined model/processor system was more likely to stimulate sleep staged as N3 than expected by chance. This was calculated by comparing the proportion of N3 in sleep with stimulation to the proportion of N3 in all sleep, as shown in Figure S3.

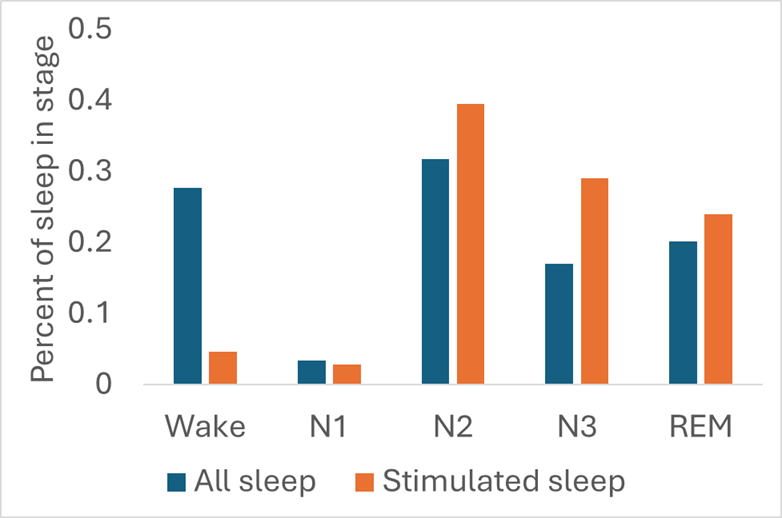

*Figure S3:* Stimulated sleep selected by the ML model+post processor is enriched for N2, N3, and REM sleep. The largest degree of enhancement was found for N3 sleep, and wake was underrepresented in the stimulated sleep.
